# Response of Soil Microbiome Structure to Biological Control Agents (BCAs) in Strawberry Greenhouse

**DOI:** 10.1101/2020.12.24.424290

**Authors:** Senlin Liu, Muzammil Hassan Khan, Zhongyuan Yuan, Sarfraz Hussain, Hui Cao, Yabo Liu

## Abstract

Continuous cropping always leads to severe abiotic and biotic problems, especially the high-intensity land utilization in greenhouses, which causes widespread concern. Effective Microorganisms (EM) and *Bacillus subtilis* (BS) have been widely used to promote plant growth and increase yields as biological control agents (BCAs). However, their effects on soil microbes are obscure. To regulate the microbial community in continuous cropping strawberry soils, we developed four soil amendments by combining EM and BS with compost. The amplicon sequencing of bacterial and fungal ribosomal markers was applied to study the response of the soil microbiome structure. We noticed a sharp increase in bacterial diversity after the addition of EM-treated high compost and BS-treated low compost, while there was no significant change in fungal diversity among treatments. Interestingly, both the relative abundance and FUNGuild predictions was consistent in revealing that BCAs may inhibit fungal pathogens in soils. Correlation analysis indicated that soil microbial community was indirectly driven by soil properties. Co-occurrence networks demonstrated that BCAs could be microecologically homogeneous through enhancing bacterial network complexity and modularity. Collectively, EM-treated high compost and BS-treated low compost can well regulate the microbial community structure and thus maintain soil health.

## 1. Introduction

Strawberries (*Fragaria× Ananassa*) are world-renowned high-value soft fruits. China’s strawberry cultivation area accounts for 40% of the world’s total in 2016. With a total production of more than $10 billion, the industry is one of the major contributors to the national economy [1, 2]. Soil microbes play essential roles in maintaining soil health and ecosystem function [3]. Long-term monocropping on the same site may cause serious problems [4]. It usually results in a dysfunctional soil microbial community structure, increased abundance of pathogenic microorganisms, and decreased abundance of beneficial microorganisms [5, 6]. For instance, over the past decade, there has been a significant decline in the richness and diversity of bacterial and fungal communities and a significant increase in the communities of *Fusarium* in the continuous cropping strawberry fields. According to recent research, the growing problem of continuous cropping in strawberry production is prevalent in all regions [3, 7].

Soil chemical fungicide is commonly used during cultivation, which may improve crop yield by killing soil pathogenic microorganisms [8]. By 2018, methyl bromide and anaerobic soil disinfestation (ASD) have blocked the spread of soil-borne plant pathogens in field settings [9] A field trial conducted by [10], showed that ASD induces changes in soil microbiome structure and strawberry disease-causing pathogens, and enhances commercial strawberry production. However, the fact that the pathogen can survive in the soil for years makes soil fungicide only partially effective [11]. Crop rotation is also known to be an option to mitigate soil pathogens. The increase in yield of corn-soybean rotation is usually attributed to microbial community in the soil, especially when it comes to disease control and nutrient availability [12, 13]. However, these traditional methods have many drawbacks, for instance environmental pollution and high costs. Consequently, we have further to address this problem with more economical and safety-friendly soil conditioners.

In general, beneficial soil microbes can compete with pathogens [13, 14]. Furthermore, these microbes help manage nutrients by making nutrients available in plants through decomposition, solubilization, iron carrier production, or symbiosis [15] A series of studies have shown that organic amendments usually have the most significant effects on microbial community in agricultural soils, such as compost or manure [16, 17]. Therefore, the use of soil amendments based on biological control agents (BCAs) and compost is considered to be a sustainable strategy[18, 19].

According to the EM Research Organization (www.emrojapan.com/how/), Effective Microorganism (EM) has been developed in Japan since the 1980s, and it has been confirmed to be composed of lactic acid bacteria, yeasts, nitrogen-fixing bacteria, and photosynthetic bacteria [20]. It has been reported that EM could increase the diversity of soil microbes and control soil diseases, thus contributing to crop growth [21, 22]. LauraNey’s research showed that the combination of EM and compost could enhance the resistance of soil food webs to drought stress as well as improving N mineralization from compost manure [23]. The successful performance of EM depends on appropriate formulation techniques and ingredients (nutrients, adhesives) for improving its durability and reliability under current environmental conditions [24]. As another BCA in soil amendment used, *Bacillus subtilis*, scientists have found that it has a good inhibition effect on a variety of plant pathogens [25], including *Verticillium sp*, *Fusarium oxysporum* and *Penicillium digitatum* [26, 27]. Besides, scientists studied the impact of the incorporation of *Bacillus subtilis* on the composition of bacterial and fungal communities in cucumber and rice rhizosphere. They found that it could be used as a plant protection agent that is compatible with the soil environment, depending on the soil type [7, 28].

Hitherto, numerous studies have focused on the effects of fertilization and soil management measures, including tillage [29], rotation [30], straw [31]. Besides, relevant studies on the addition of soil amendments in the strawberry field mainly gave priority to its pathogenic fungi. In contrast, there are few studies on the function and co-occurrence network of strawberry soil microbes [32, 33].

In this study, we developed a series of soil amendments by combining compost with two popular commercial BCAs (EM and BS). A field experiment was carried out in a long-term continuous cropping strawberry greenhouse in southern China, and we applied 16S rRNA amplicon sequencing [34] for further study. We hypothesize that, with soil amendments processing, the diversity and co-occurrence patterns among strawberry soil microbiome could be improved ideally. Besides, we assume that the soil ecological function has predictable heterogeneity. The objectives of our study are: (1) to regulate the soil microbial community structure and control soil pathogens in continuous cropping field; and (2) to put forward a theoretical and practical basis for the sustainable production of strawberry and other plants from the perspective of microbial ecology.

## 2. Materials and methods

### 2.1. Soil amendments preparation

The soil amendments compared were organic compost with two biological control agents (BCAs): Effective Microorganisms (EM) and *Bacillus subtilis* (BS). Rice bran and soybean meal were blended clinched alongside a ratio of 1:2 (dry weight) to serve compost for the processing of soil amendments [35], and the detailed parameters for compost are provided in the supplement TableThe EM (~1 × 10^9^CFU/mL) and BS (~1 × 10^11^CFU/mL), are produced by Jiangsu Warner Biotechnology Co., Ltd. The main components of EM were *lactobacillus plantarum*, *Lactobacillus acidophilus*, *Lactobacillus pentose*, yeast, *Bacillus pumilus*, nitrifying bacteria and metabolites. The main component of BS is *bacillus subtilis*. The original EM agent is of pH 3.5 compared to 7.0 for BS. EM was activated by using 1.0 L mother culture EM • 1® and mixed with 500 mL unsulfured molasses in bioreactor under anaerobic conditions [36]. After a week, the activated EM with the pH value above 4.0 could be accessible, and BS can be used directly. The two sorts of BCAs were diluted with chlorine-free water at a proportion of 50 times, and the compost was produced with the application of EM, BS at turning. In order to optimise the proportion of compost in the soil amendments, we have set two levels of compost content in each BCA.

As stated by those extent for unit zone utilized within greenhouse, four soil amendments were prepared, which were: (I) EM 1ml/m^2^+compost 125g/m^2^ (EM1), 1.05kg per treatment repetition; (II) EM 1mL/m^2^+compost 250g/m^2^ (EM2), 1.8kg per treatment repetition; (III) BS agent 1ml /m^2^+compost 125g/m^2^ (BS1), 1.05kg per treatment repetition; and (IV) BS agent 1ml/m^2^+compost 250g/m^2^ (BS2), 1.8kg per treatment repetition. It is worth noting that EM and BS are calculated based on the original concentration. Following preparing, the soil amendments were stored and then utilized within greenhouse experiments.

### 2.2. Experimental design and Sampling

Our greenhouse experiment and design were carried out in Baitu Town, Zhenjiang City, Jiangsu Province, China (31°57’N, 120°09’E). The region has a Northern Subtropical climate, with average annual precipitation of 1022 mm and mean yearly temperature of 17.1°C [1].

The plots in the greenhouse are arranged in a randomised block design with three replications per treatment, and no application of soil amendments as the control. The specific treatments were control, EM1, EM2, BS1 and BS2, respectively. The greenhouse area was 10 m wide and 60 m long, which contained five experimental plots for five treatments, and each plot in the greenhouse is 0.5 m wide and 12 m long. Strawberries are planted in double rows with 20cm interval, and the soil type is loam according to Soil Classification Retrieval System of China.

This greenhouse has been used to plant strawberry for more than 5 consecutive years prior to this experiment, and we have witnessed a decline in strawberry production and growth in recent years. As a traditional method of alleviating continuous cropping pathogens, the greenhouse was closed in July 2018 to make use of sunlight and weeds were removed in August. Strawberries were transplanted in September, while soil amendments were introduced to the soil layer and covered with an agricultural plastic film according to the treatment process detailed in 2.1. Water was conveyed through drip irrigation and maintained under the same agricultural management model (Supplementary Fig 1). It was guaranteed the strawberries cultivated in greenhouse belonged to the same variety (*Benihoppe*).

**Fig 1.**
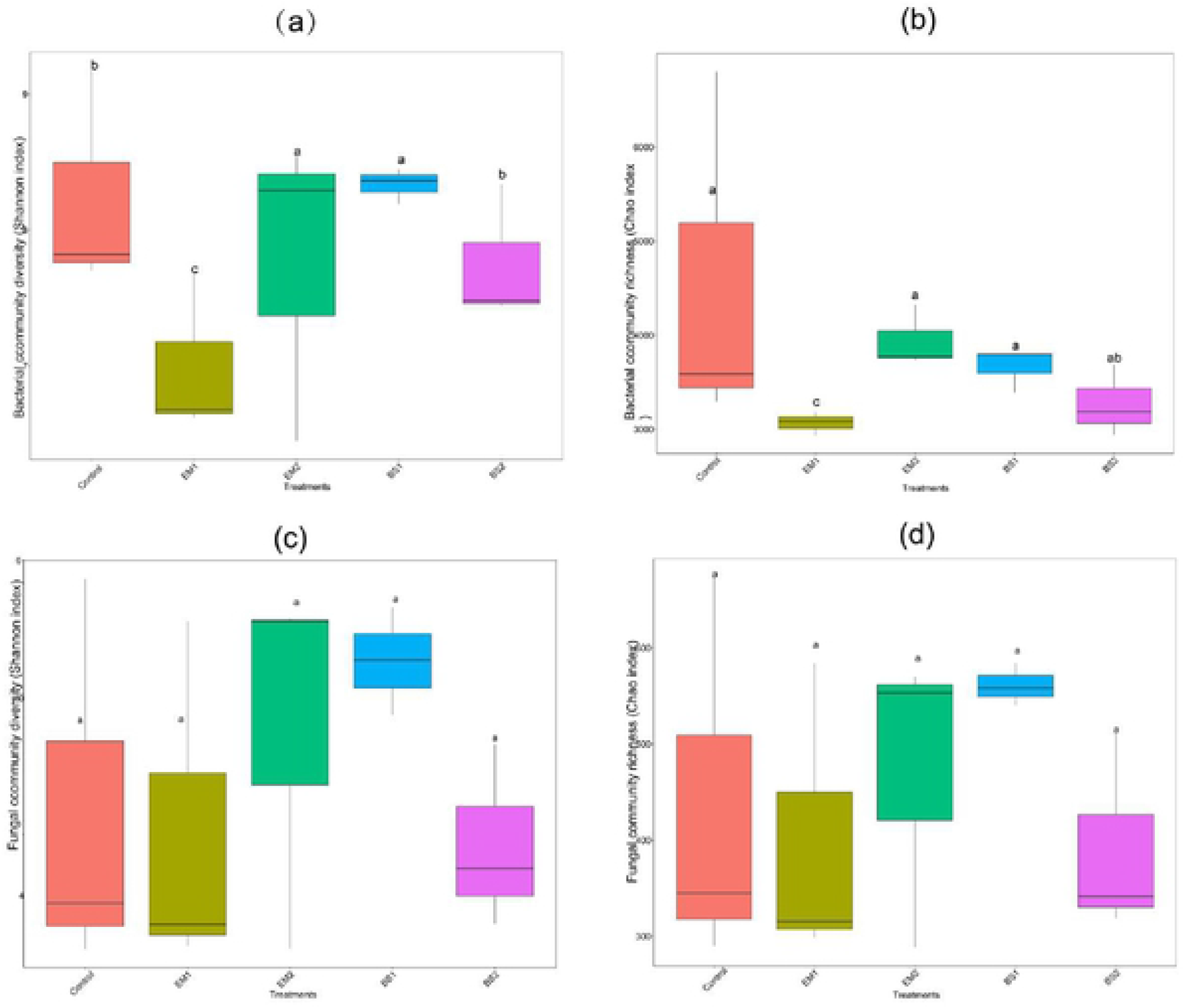
Comparative analysis of the alpha diversity index in different treated soils: (a) Shannon of bacterial 16S rRNA gene, (b) Chao of bacterial 16S rRNA gene, (c) Shannon of fungal ITS gene, (d) Chao of fungal ITS gene, were calculated by five treatments. Statistically significant differences were dctcnnincd by one-way ANOVA (P < 0.05).

Three single-well soil samples were collected randomly from each treatment plot as three replicates. greenhouse soils (0 - 20 cm depth) were gathered from the greenhouse with an S-pattern in December 2018, when the strawberries were in the fruiting stage. Then afterward collecting the strawberry soil, a total of 15 fresh soil samples were put in aseptic plastic bags and brought back to the laboratory for storage at 4°C and −80°C, separately.

### 2.3 Analysis of soil physicochemical properties

Soil pH (soil:water=1:5, w/v) was determined using a pH meter with a glass electrode (FE20-Five Easy Plus ™, Switzerland) [37]. Total organic C(TOC) was determined according to the vitriol acid-potassium dichromate oxidation method [38]. Total nitrogen (TN) was measured based on direct combustion using an elemental analyzer[39]. C/N ratios were measured by the ratio of TOC to TN. Inorganic N (NH_4_^+^-N and NO_3_^−^-N) of soil was drawn with 2 mol/L KCl (soil:KCl=1:10, w/v) by shaking (1h, 200rpm) and filtering through polysulfone membrane, before colorimetric determination requiring a continuous-flow analyzer [40]. Available K (AK) and total K (TK) of the soil samples were identified using flame photometry method [41], and available P (AP)was tested by the molybdenum blue method. Soil physicochemical properties are presented in Supplemental Table 1.

### 2.4. DNA extraction and 16S rRNA and ITS amplicon sequencing

In order to ensure the validity of the experiment, the soil was kept in the refrigerator for only one week before the soil DNA was extracted. The microbial DNA of fifteen soil samples was extracted from 1.0 g of each sample by the E.Z.N.A.® Soil DNA Kit (Omega Bio-tek, Norcross, GA, U.S.) conforming to the manufacturer’s instructions. The V3-V4 region of the 16S rRNA gene and the ITS1 region of the fungal ITS gene were selected as specific fragments for detection of bacteria and fungi using primers 338F/806R [42] and ITS1F/ITS2 [43], respectively. PCR reactions were performed in triplicate 30 μ l mixtures containing 10 ng of template DNA, Phusion® High-Fidelity PCR Master Mix (New England Biolabs) 15 μl, 2μmol/L Primer 3μl. The PCR reactions for the 16S V3-V4 rRNA gene were conducted following the process: initial denaturation under 95°C for 3 min, 30 cycles consisting of denaturation for 30s at 95°C, annealing at 56°Cfor 30 s, followed by 72 °C for 45 s, and a final extension for 5 min at 72°C; as for the ITS gene, the following procedure was followed: an initial denaturation step at 95°C for 3 min, followed by 35 cycles at 94°C for 30 s, 55°C for 30 s and 72°C for 45 s, and finally an extension of 10 min at 72 °C [44]. The resulted PCR products were extracted from a 2% agarose gel and further purified with GeneJET TM Gel Extraction Kit (Thermo Scientific) as stated by the protocol of manufacturer. The library quality was assessed by the Qubit@ 2.0 Fluorometer (Thermo Scientific).

Single-end of 16S rRNA gene and ITS1 sequenced on an Ion S5TM XL platform (Wang et al., 2018) by Novogene Genomics Institute (Beijing, China). Those raw reads were deposited into the NCBI Sequence Read Archive (SRA) database (Accession Number: SUB7456591).

### 2.5. Bioinformatic processing and Analysis

The naive-Bayes, BLAST+-based, and VSEARCH-based classifiers implemented in QIIME (V2.0, http://qiime.org/) [45] designed for classification of bacterial 16S rRNA and fungal ITS marker-gene sequences that were evaluated in this study. Then, sequences were quality controlled (> 25 score and the length of 200 bp), and according to the corresponding barcode assigned to different samples. Sequences with ≥ 97% similarity were assigned to the same operational taxonomic units (OTUs) [46], then the bacterial OTUs of the representative sequences were performed by the Silva (Version 132) database (https://www.arb-silva.de/) [47]. The Heatmap, Barchart and correlation analysis (RDA, CCA) were displayed with R-Studio (Version 3.6). We defined specific OTUs as “abundant” when their average relative abundances were above 0.05% across all samples following [48]. For the Mantel test, it focused those soil physicochemical properties that significantly correlated with abundant OTUs by the Bray-Curtis dissimilarity algorithm. The differences between treatments were analysed by one-way ANOVA (P < 0.05) using the SPSS 25.0 software.

FAPROTAX (version 1.1) [49] was employed to annotate the functional annotation of bacterial community in the normalized OTU TableFAPROTAX (http://mem.rcees.ac.cn:8080/root) is a manually constructed database that maps prokaryote to possible ecological functions (nitrification, denitrification or fermentation) or metabolic. For instance, if all cultured strains of the bacteria have been identified as nitrification types, FAPROTAX assumes that all uncultured genera are the same functional group.[50]. Correspondingly, FUNGuild is an ITS-based functional prediction software launched in 2016, and is currently based on a classification prediction called ‘guild’, which is based on data integrated from published literature [51]. There are 12 categories of pathogenic bacteria, animal pathogens and wood decay fungi.

### 2.6. Co-occurrence network analyses

In order to illustrate the co-occurrence interaction between bacteria in strawberry greenhouses, network analysis was performed on the abundance of the top 80 genera between treatments. We adopted Spearman’s correlation to obtain the strong correlation (r> | 0.8 |) and significant correlation (P <0.05) between taxa. Next, we used Cytoscape version 3.8.0 [52] to visualize the network structure. The size of each node stands for relative abundance of the genus of microbe. The colour of each node was distinguished depending on the level of phylum. Correlation was shown as an edge (positive correlation = grey; Negative correlation = red); At the same time, Gephi (v.0.9.2) and Network Analyzer were utilized to calculate the obtained network topology parameters(number of nodes and links, network density, shortest paths, network diameter, average neighbors, and clustering coefficient) to represent the co-occurrence relationship between genera [53].

## 3. Results

### 3.1. Richness and diversity of bacterial and fungal communities

From 15 soil samples, we obtained a total of 1,537,746 high-quality V3-V4 sequences of 16S rRNA and 1,202,670 high quality ITS1 sequences, average read length of bacteria and fungi were 437 and 282 bp, respectively. The sequences were grouped into 3888 bacterial operational taxonomic units (OTUs) and 1234 fungal OTUs at 97% sequence similarity cutoff (Supplementary data 1). For the α-diversity, indexes including observed OTUs, Chao1, ACE, Shannon, and Simpson of bacterial and fungal communities were observed (Supplemental Table 2). All coverage of soil bacteria and fungi was more than 97.9%, indicating the current sequencing depth in this study was accurate and reliable. The Shannon index (Fig 1a) showed that the bacterial diversity of the EM2 and BS1 treatments was significantly higher than that of the control (P < 0.01); the Chao index (Fig 1b) showed that the bacterial richness of EM1 was significantly lower than that in the control (P < 0.05). Conversely, we did not see significant differences in community richness and soil fungi diversity (Fig 1c, d).

The Venn diagram shows that the distribution of OTUs in the microbial community varied among the different treatments (Fig 2). A total of 1314 OTUs were shared among the five soil treatments, accounting for 33.80% of the total. In addition, 311, 57, 90, 107 and 282 OTUs were unique in the control, EM1, EM2, BS1 and BS2 treatments, respectively (Fig 2a). Interestingly, in the same BCA, the number of bacterial unique OTUs in the higher composts increased significantly. Soil fungi shared 323 OTUs among the five treatments, accounting for 26.16% of the total OTUs. In the order of the above soil treatments, there were 129, 45, 25, 66 and 20 unique OTUs for soil fungus, respectively. (Fig 2b). We found a significant reduction in the number of unique OTUs in compost with the same agent.

**Fig 2.**
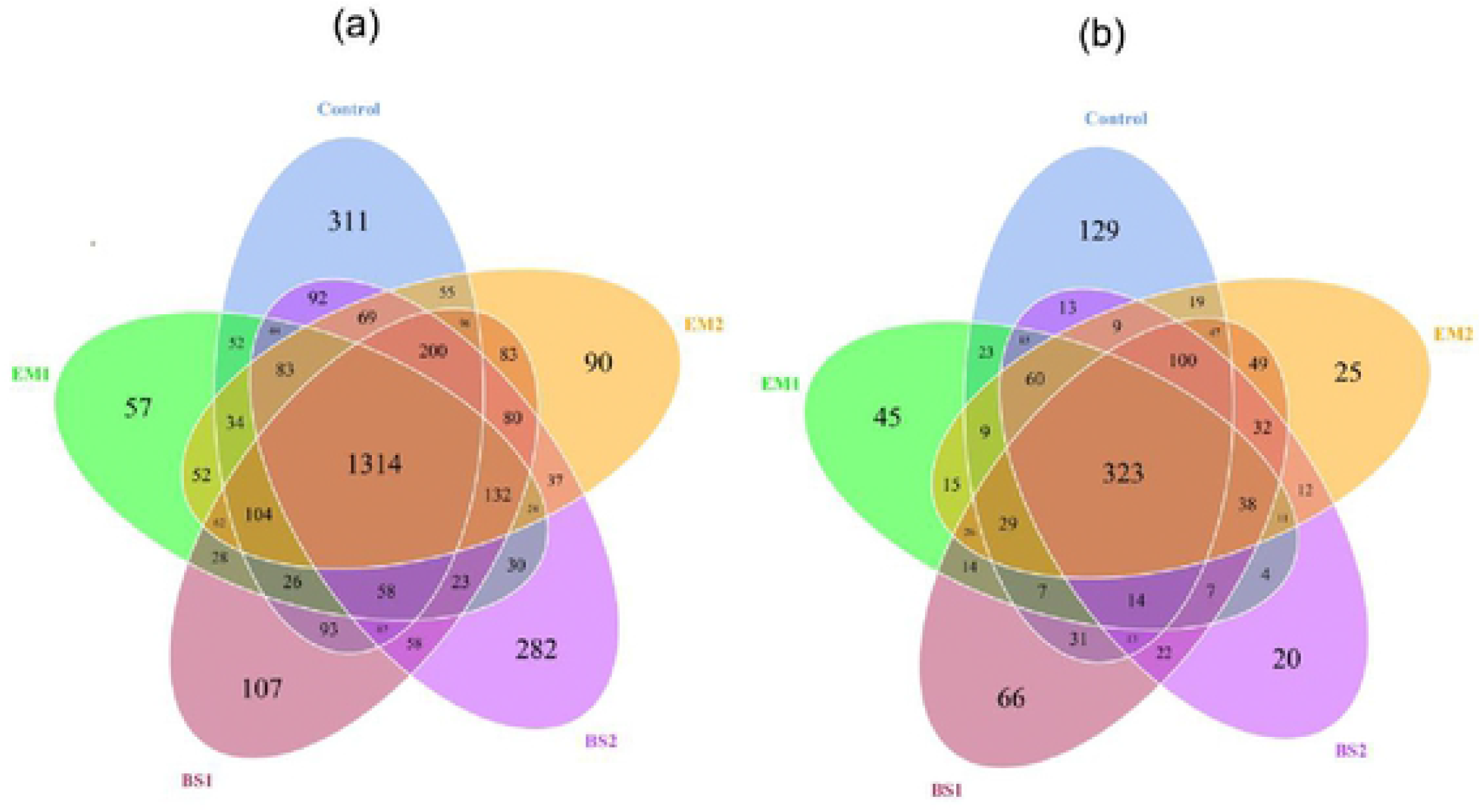
Venn diagram showing the unique and shared bacterial OTUs (3% distance level) among the different libraries in CK (pink), EMI (green), EM2(blue), BSl(red) and 8S2 (yellow) treatments: (a) Venn diagram of bacterial OTUs between five treatments; (b) Venn diagram of fungal OTUs between five treatments. The numbers in one circle denote unique OTUs, and numbers in l\vo or more intersecting circles denoteshared OTUs.

### 3.2. The core microbiome at genus level

According to the taxonomic identification, there were 42 phyla, 50 classes, 114 orders, 223 families, and 570 genera in bacterial community, while fungal OTUs could be classified into 10 phyla, 29 orders, 62 orders, 101 families and 144 genera. Nineteen bacterial genera with relative abundance greater than 1% were identified (Fig 3a), and the most abundant genera are *Rhodanobacter* (8% - 21%), *Bacillus* (3% - 8%), *Arachidicoccus* (1% - 6%). *Rhodanobacter* (p_Proteobacteria)*, Bacillus, Arachidicoccus, Thermoflavifilum, Sphingomonas* (p_Proteobacteria) *, Pseudomonas* (p_Proteobacteria)*, and Streptomyces* (p_Proteobacteria). Among them, there were significant differences among different treatments in phyla Firmicutes and Bacteroides belonging to Proteobacteria (P < 0.05). All four soil amendment treatments increased the total relative abundance of dominant genera. Among them, EM1 was the highest (52.2%), and the control greenhouse was the lowest (15.83%). Compared with the control treatment, the application of EM1, EM2, and BS2 raised the relative abundance of *Rhodanobacter* and *Arachidicoccu*. However, there was no significant difference between two compost treatments. At the same time, EM1, EM2, and BS2 significantly reduced the relative abundance of *Sphingomonas* in soil. Both BS1 and BS2 treatments significantly advanced the relative abundance of *Bacillus*. All the four soil amendment treatments remarkably reduced the relative abundance of *Thermoflavifilum*, with the most significant decline in EM2.

**Fig 3.**
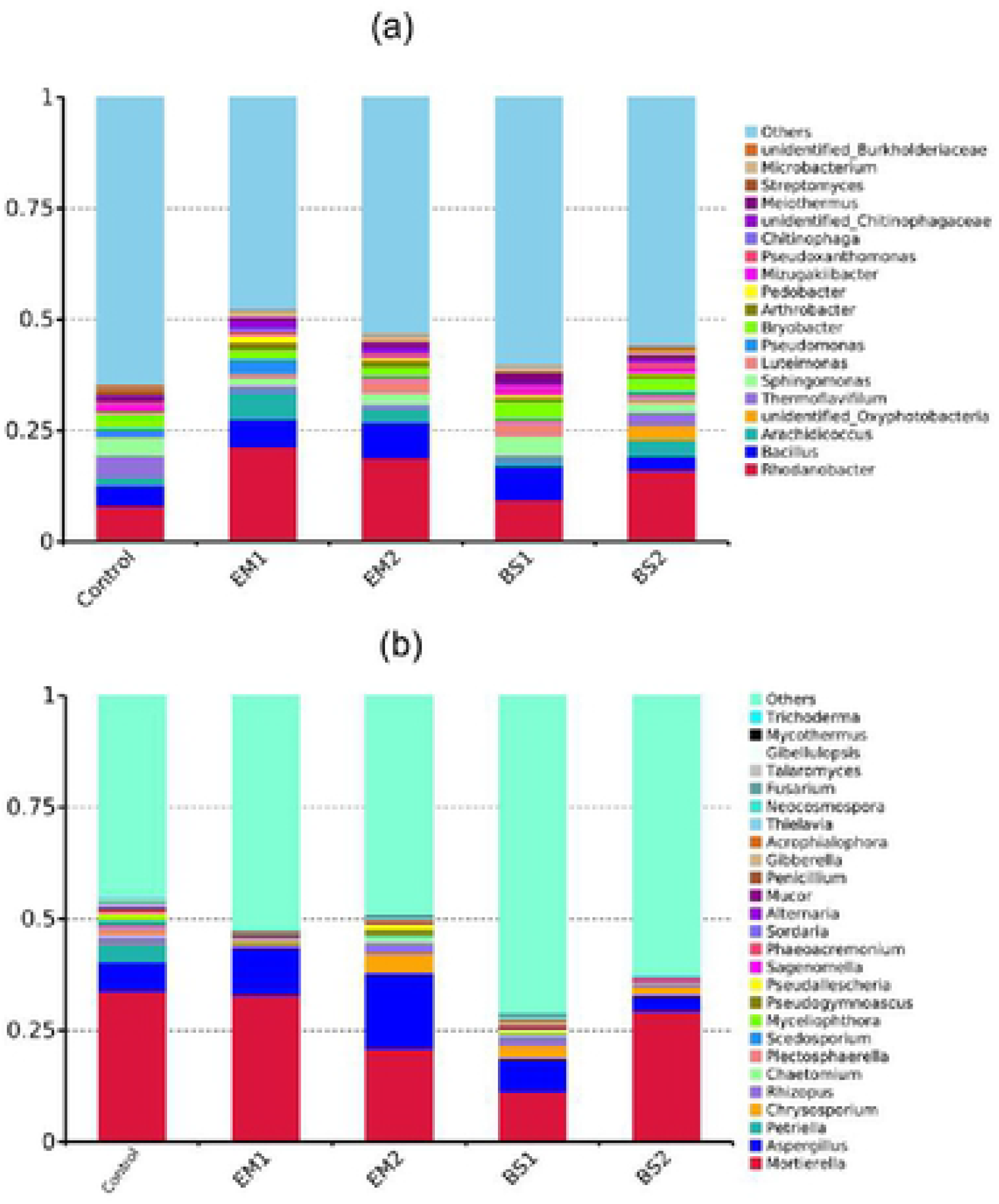
Changes in the relative abundances of bacterial (a) and fungal (b) dominant genera(b) under different treatments of strawberry soil, proportional distribution of taxa with abundance >1%. Different letters represent statistical significance at P <0.05.

Considering that fungi are key microbes for soil-borne diseases, it is necessary to focus our attention on them. After a previous classification in the literature, we identified 9 genera of plant pathogenic fungi in the strawberry continuous cropping soil (Supplemental Table 3), of which the first 7 genera (including *Aspergillus*, *Rhizopus* and *Penicillium*, etc.) are dominant genera (Fig 3b) and the remaining 2 genera are rare taxa. The average total abundance of these pathogens was 9.94% (CK), 0.98% (EM1), 8.27% (EM2), 3.60% (BS1) and 0.93% (BS2), respectively. The total abundance of these pathogenic fungi decreased with the application of the soil amendments, with EM1, BS2 decreasing the most. Further ANOVA analysis revealed that the genera *Aspergillus*, *Rhizopus*, *Penicillium*, *Fusarium*, *Alternaria*, *Mucor* and *Botrytis* differed significantly among different treatments. In particular after the implementation of soil amendments, the relative abundance of *Rhizopus*, *Penicillium* and *Fusarium* all decreased significantly (p < 0.05).

### 3.3 Correlations of abundant microbial taxa with edaphic variables of soil Characteristics

We carried out the correlation analysis using the soil physicochemical properties as well as the selected abundant OTUs of sequences (Fig 4a). In the entire dataset, there exsisted In the whole data set, the abundant taxa contained 159 bacterial OTUs and 28 fungal OTUs (Purple dots in figure 4). CCA indicated that 54.43% of the total variance within abundant bacterial taxa was explained by the first (29.79%) and second (24.633%) ordination axes (Fig 4a). The variation in bacterial composition was significantly explained by TN, AP, AK and C/N ratios. The EM2, BS1 treated samples occupied a richer bacterial community, which is consistent with the previous findings. The richer bacterial taxa in the EM1 treated samples were closest to TN, AP, AK, and they correlated well with each other. For the fungal community (Fig 4b), RDA indicated that 82.293% of the total variance within abundant bacterial taxa was explained two ordination axes. There is a clear distinction between the different treatments. However, the correlation between the abundant fungal taxa in the different treated soils and soil physicochemical properties was weak.

**Fig 4.**
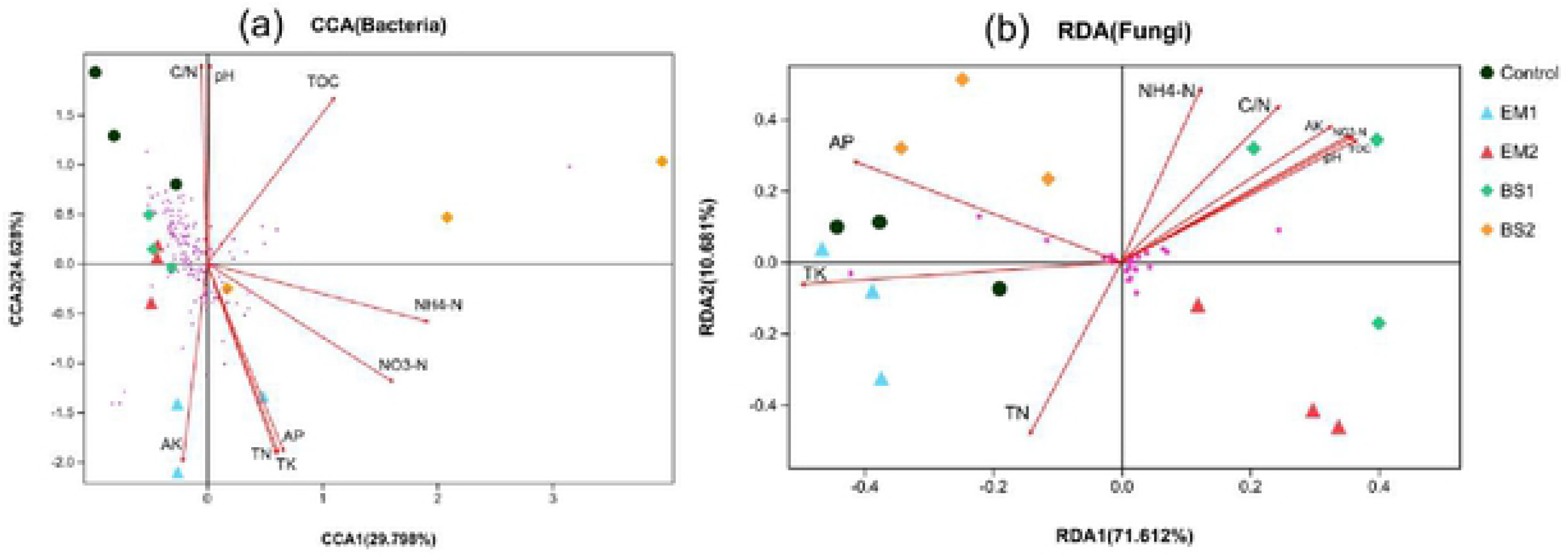
RDA and CCA demonstrating the relationships between soil environmental factors and soil microbial communities(bacterial (a), fungal (b)) after application of soil amendments. The soil microbial communities selected the abundant OTUs represented by more than 0.5% relative abundance. The length of each arrow indicates thecontribution of the corresponding parameters lo the structural variation. The treatments arc indicated in different colors respectively. Soil factors indicated in blue text include total carbon (TOC), total nitrogen (TN), total phosphorus(TP), NH_4_^+^-N, N0_3_^−^-N, pH, Total polassium(TK), Available potassium(AK), Available phosphorus(AP).

Mantel test (Supplementary Table 4) revealed significant (P◻values based on 999 permutations) relationships between bacterial abundant community and TN, AP, AK and C/N ratios. On the other hand, there were significant relationships between fungal abundance communities and the presence of AP and TK only.

### 3.4. Potential roles of key microbial players in the strawberry soil with amendments

The FAPROTAX database has been extensively used to analyse the biogeochemical cycling processes of bacterial communities. We assigned 768 out of 3,863 bacterial OTUs (19.9%) to at least one microbial functional group. Sixty-seven predicted functions were identified. Then the most abundant 25 functional groups were further evaluated for their relative profiles in the different soil samples (Fig 5a). Chemoheterotrophy and aerobic_chemoheterotrophy were the two highest relative abundance in different treatments among the putative functions, accounting for 15.55% and 14.90% of the total respectively. Many functional groups in the soil were more abundant after BS1, BS2 treatment and were reduced after EM1, EM2 treatment. Among them, BS2 application significantly lowered the relative abundance of chemoheterotrophy, aerobic_chemoheterotrophy function. Whereas EM1, EM2 application remarkably reduced nitrogen_respiration and nitrate_respiration function. After the four soil amendments applications, there was a significant decline in the function of predatory_or_exoparasitic, invertebrate_parasites.

**Fig 5.**
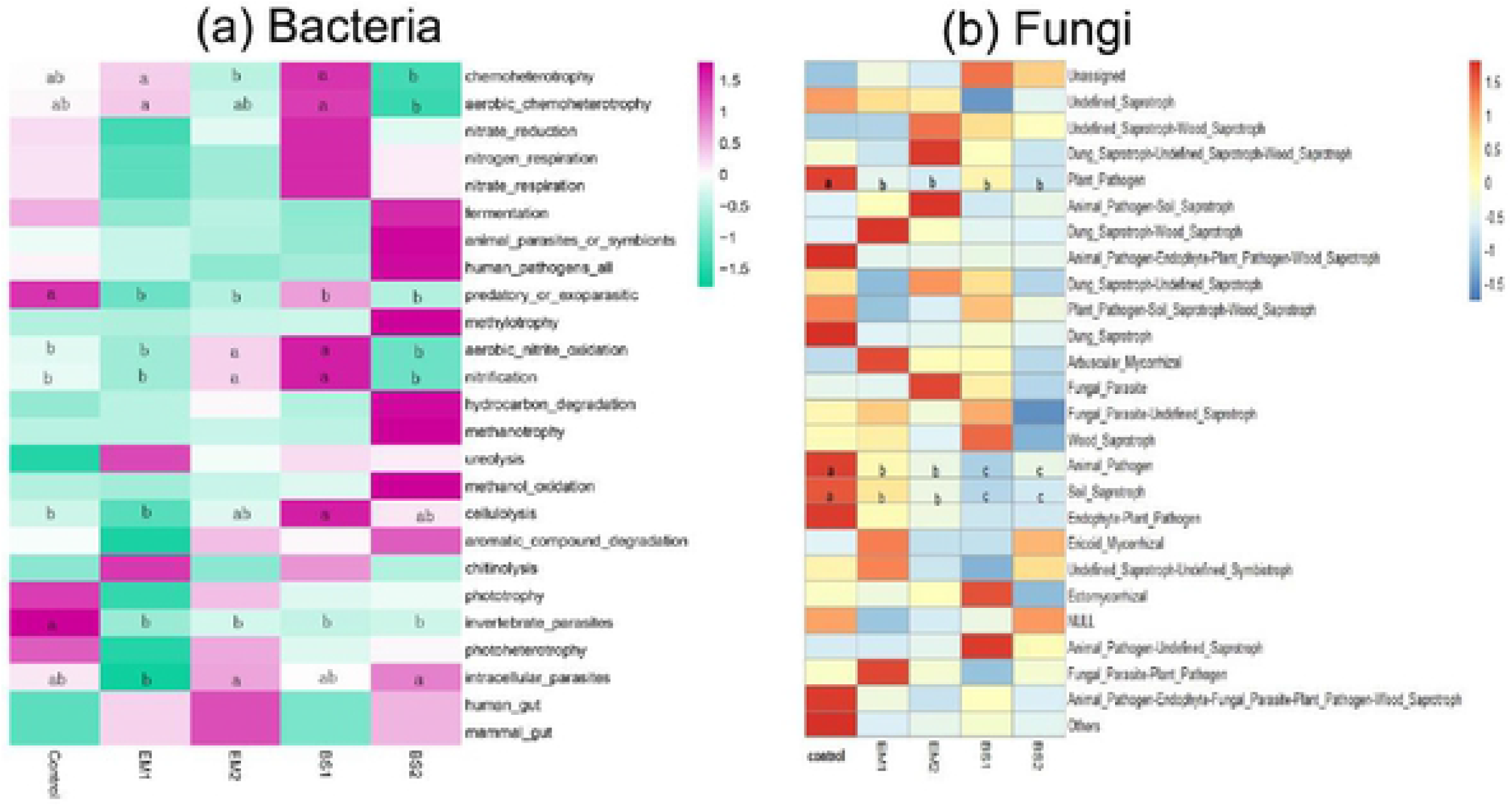
Healmap showing relative functional abundance predictions of the bacterial communities based on FAPROTAX (a), and fungal communities based on FUNGuild (b). The color code represents the row z-score. Different letters (a, b, ab, c) represent statistical significance al P <0.05.

The function of fungal microbial community was predicted by FUNGuild. A total of 825 OTUs were classified into fungal guilds, accounting for 66.86% of all OTUs. As shown in Fig 5b, Unassigned and Undefined_Saprotroph functional groups of fungi dominate the top 25 functional guilds, with average abundances of 54.27% and 40.45% respectively, while other functions are less predominant (about 6%). All soil amendments significantly reduced Plant_Pathogen, Animal_Pathogen and Soil_Saprotroph functional fungi. Specifically, for the latter two functions, soil amendments with BS was more significant than with EM applications.

### 3.5. Structure and composition of bacterial co-occurrence networks

Co-occurrence network analysis was conducted to assess the complexity of the interactions among bacterial genera detected in strawberry soils treated with different amendments. Spearman was used to calculate the correlation between the top 80 bacterial genera in the soil. Then, we selected the Cytoscape software to visualize the co-occurrence network (Fig 6) and evaluate several vital topological properties (Table1).

**Table. 1.**
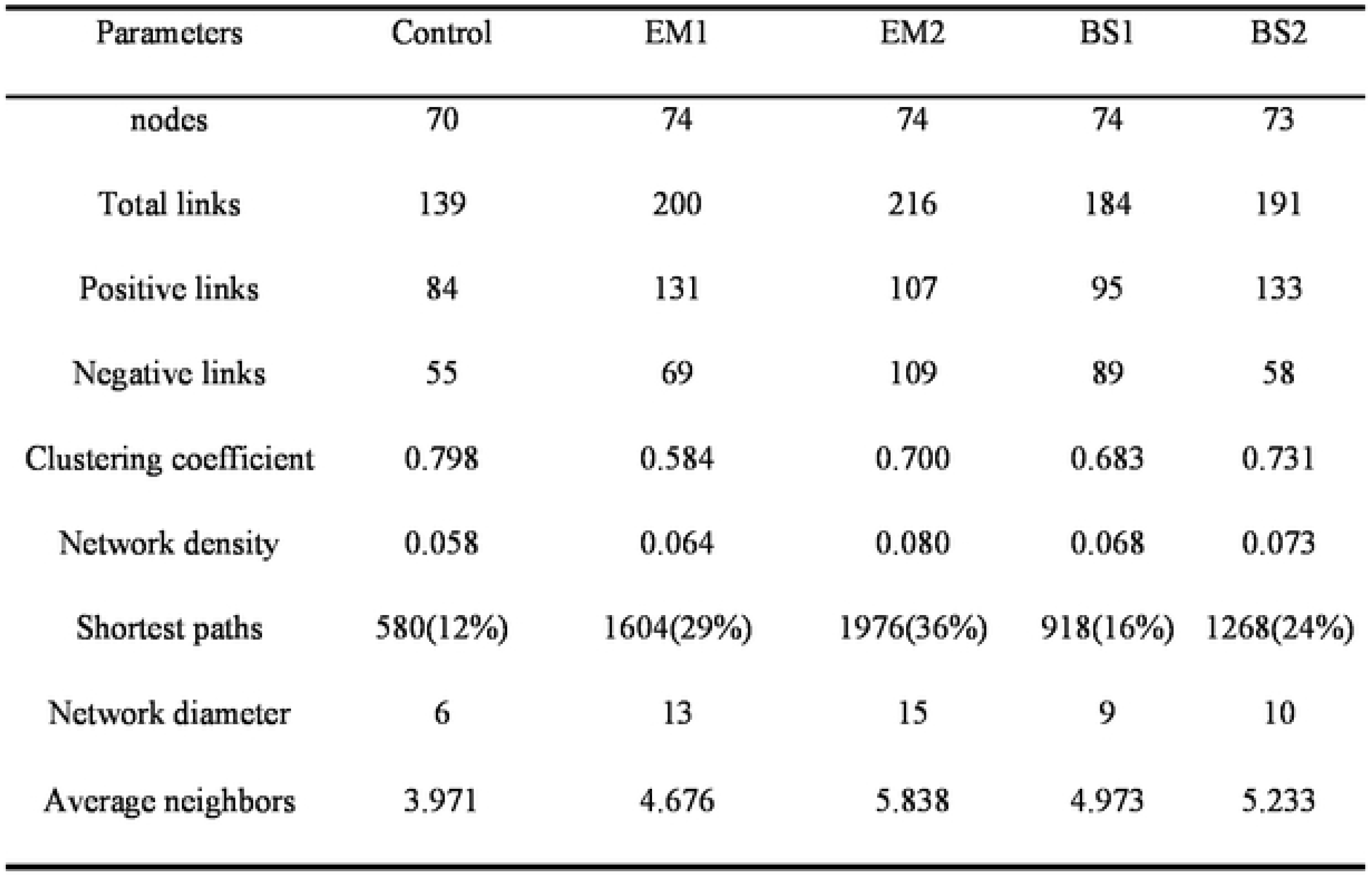
Topological properties of of Correlation network diagram of soil bacterial communities at genus level in different treatments.

**Fig 6.**
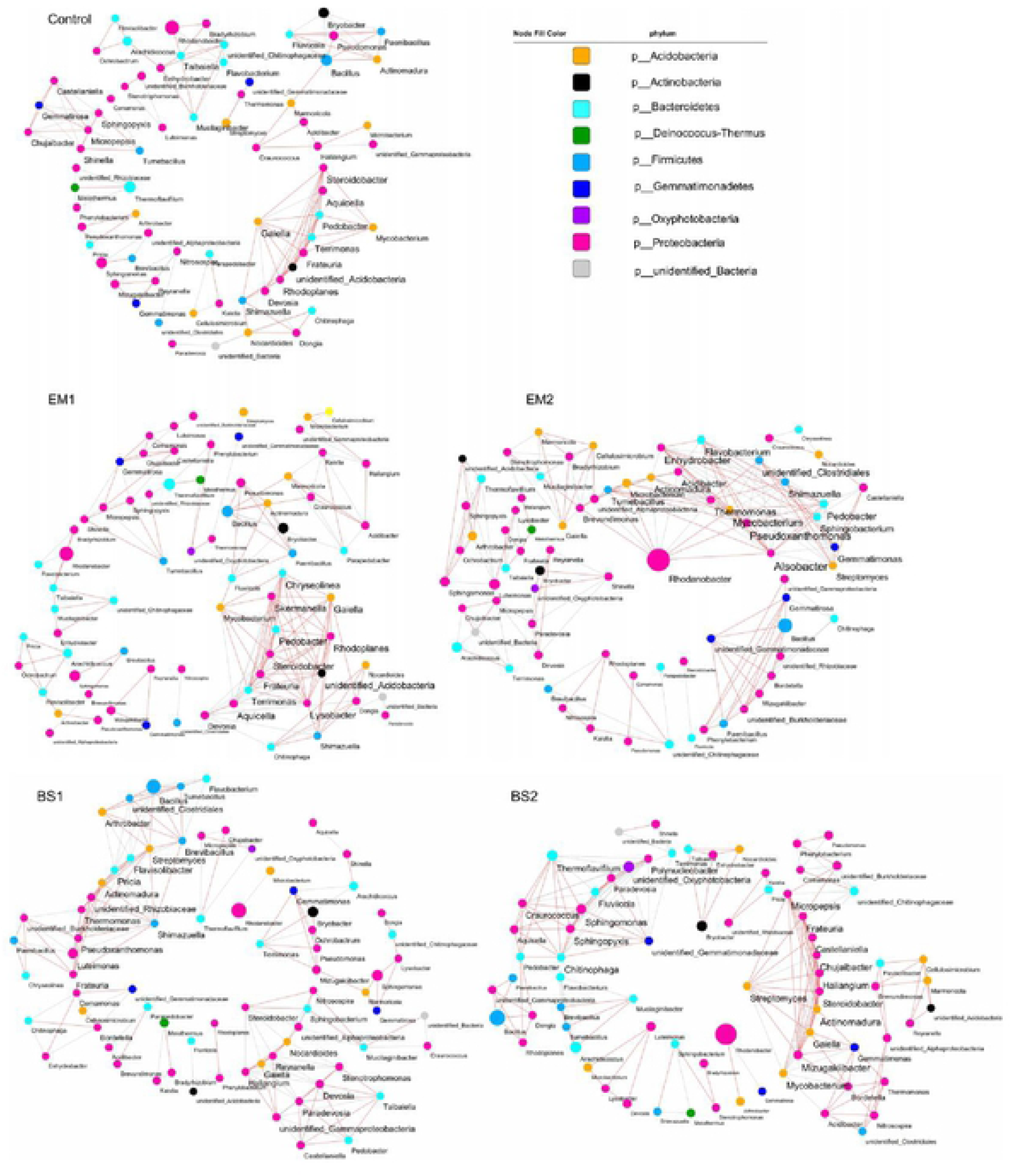
Co-occurrence network diagram of soil bacterial communities at genus level between different treatments. Based on Spearman correlation, Cytoscape was used to construct bacterial co-occurrence network.Correlation is shown as edge (positive correlation = gray;Negative correlation = light red), correlation coefficient r> |0.8|, and P <005. The size of nodes is positively correlated with relative abundance of genus, and !he color of nodes is distinguished by phylum level.

The bacterial network was composed of nodes and edges, and there were 70-74 nodes with significant correlations. The results showed that nodes, total links, positive links, negative links, network density, network diameter and average neighbors all increased after the addition of soil amendments. We easily found that the bacterial network of EM2 was the most complex and compact, and it held the highest topological properties of total links (216) and network density (0.080); however, the simplest network (control) was only 139 and 0.058, respectively. Moreover, total bacterial links and network diameter in soil amendment with EM, were all larger than with BS, and had greater network complexity. Compared with EM1, the soil bacterial network was more balanced with EM added compost (positive links are more similar to negative links). Compared with BS1, positive links of soil bacteria in BS2 increased greatly, while negative links decreased.

Utilizing the degree of connectivity between microbes, we sought to find the keystone genera within each network. The higher the degree index, the closer the relationship between the genus and other taxa. The degree value of *Gaiella* (p_Acidobacteria), *Rhodoplanes* (P_Proteobacteria) and *Steroidobacter* (P_Proteobacteria) in control soil were the highest, but they were all lower than those in other soil amendments. In EM1 soil, the degree values of *Lysobacter* (P_Proteobacteria), *Aquicella* (P_Proteobacteria) and *Rhodoplanes* (P_Proteobacteria) were the highest. In EM2 soil, the degree values of, *Alsobacter* (P_Proteobacteria), *Pseudoxanthomonas* (P_Proteobacteria) and *Enhydrobacter* (P_Proteobacteria), *Flavobacterium* (P_Bacteroidetes), *Rhodanobacter* (P_Proteobacteria) were the greatest, and it was noted that the relative abundance of *Rhodanobacter* was also the richest in the community. In BS1 soil, the degree values of, *Patricia* (P_Bacteroidetes) and *Flavobacterium* (P_Bacteroidetes), *Shimazuella* (P_Firmicutes) were the highest. In BS2 soil, the genera with the highest degree were *Mizugakiibacter* (P_Proteobacteria) and *Actinomadura* (P_Actinobacteria), *Chitinophaga* (P_Bacteroidetes).

## 4. Discussion

In this study, by using amendments synthesized by BCAs and compost, we have revealed details of the soil bacterial and fungal community structure through high-throughput sequencing. The diversity of microbial community is an indicator of the effectiveness of agricultural practices [54]. Although the succession pattern of bacteria and fungi in continuous cropping strawberry fields remains unclear, the degradation of soil can be partly explained by changes in the diversity and structure of the microbial community. Some researchers found that the richness and diversity of bacterial and fungal community would be greatly reduced with the increase of continuous cropping years (especially over five years) [44]. By adopting different soil amendments, EM2 and BS1 could promote bacterial diversity, while EM1 can significantly reduce soil bacterial richness. In addition, the alpha-diversity indices of soil fungal community did not change significantly among treatments. Previous studies have shown that EM application improved soil microbial diversity [55, 56]. This is similar to the results of the EM2 treatment in our study. It may be due to the fact that EM bacterial agent is suitable for a higher proportion of compost, and the number of unique OTUs of bacteria with higher compost increased significantly in the same BCAs (Fig 2a). Recent studies have shown that the application of BCAs of *Bacillus*-based formulates does not decreased the total microbial diversity and community [57]. Instead, *Bacillus subtilis* increased the bacterial diversity in tobacco rhizosphere soil [58], which is also verified by our findings.

The structures of microbial community have undergone profound changes. Proteobacteria were the most abundant bacteria associated with disease inhibition in the soil with long-term monostrophic fertilization [6]. In our study, most of the dominant bacteria with significant differences (P < 0.05) belong to Proteobacteria whose abundance increased to different degrees after adding soil amendments. This may be one of the factors that guarantee strawberry soil health. We illustrated that when soil amendments were introduced, the total relative abundance of dominant bacterial genera increases significantly from 15.83% (control) to 52.2% (EM1). However, among the genera of soil fungi (Supplemental Table 3), the total relative abundance of seven pathogens decreased with the application of soil amendments, with the greatest decrease in EM1, BS2. The relative abundance of *Fusarium*, the most well-known of the soil and plant pathogens, decreased significantly (p < 0.05) after adding soil amendments. Previous studies have shown that the application of BCAs and certain organic matter can effectively inhibit soil pathogens, including *Verticillium* sp, *Fusarium oxysporum* and *Penicillium digitatum* [59, 60]. Besides, it has been shown that *Bacillus* and *Trichoderma* (components of EM) can protect host plants against pathogens [61, 62]. In this study, the relative abundance of the corresponding microbial population (especially Bacillus) was significantly increased by adopting BS-based soil amendments. This suggests that soil amendments may lead to increased competition for resources and antagonistic between bacteria and pathogens in composting soils. Therefore, through positive interaction, the indigenous microbial population benefits from introducing microbes into soil systems.

The microbial community consists of a large number of abundant and rare taxa. In most ecosystems, the abundance of microbes contributes to microbial biomass and mineralization of organic matter [63]. In agricultural soils, microbial communities are affected by multiple factors such as sampling time, carbon and nitrogen sources, soil water content and plant physiological status [64]. These factors may be related to microbial community assembly. Both the dominant bacterial and fungal taxa were explained by environmental factor correlation analysis, indicating their high correlation with key soil physicochemical properties. Consistent with other results, our results showed that EM2, BS1 treatments occupied a richer bacterial community, and there is significant correlation between the bacterial community and TN, AP, AK and C/N ratios, whereas the fungal community was only significantly correlated with AP and TK. Thus, the application of four soil amendments reconstructed soil microbial communities through changes in soil physicochemical properties.

Studies have shown that green manure of soybean promoted the increase of functional bacteria like nitrogen-fixing bacteria, nitrifying bacteria and denitrifying bacteria in soil, indicating that green fertilizer application promoted the nitrogen fixation and nitrogen cycle process in soil [50, 65]. Based on FAPROTAX function prediction, we estimated that the BS-based soil amendments promoted multiple functions of soil bacteria, such as the aerobic nitrite oxidation, nitrification and cellulolysis. Nevertheless, the EM-based soil amendments significantly reduced multiple functions of soil bacteria. Accordingly, FUNGuild prediction showed that soil amendments significantly reduced the taxa of Plant_Pathogen, Animal_Pathogen and Soil_Saprotroph functional fungi, especially the decrease of plant_pathogenic functions matched the decrease of these fungal pathogenic taxa shown in table 1 above. Consistently, BS containing soil amendments showed more significant inhibition against these harmful pathogens than EM-containing soil amendments.

Further analysis of the co-occurrence network of 80 dominant genera in soil microbial community showed that the interaction among bacteria in strawberry soil after applying soil amendments was more complicated than that in the control soil. In addition, the total bacterial links of EM (EM1, EM2) were higher than those of BS (BS1, BS2). The bacterial network of EM2-treated soils was the most balanced and complex. Based on these results, we assumed that the application of EM and more compost in strawberry soils made bacterial communities more complex and modular, which made it easier for specific bacteria to establish symbiotes in agricultural soils [66]. According to this hypothesis, the colonization rate of relatively single flora in BS-treated soil was lower than that of mixed flora EM, which maintained the health and balance of soil microbes weakly. In the bacterial network of different treatments, the keystone genera have undergone significant changes, but they all generally belong to Proteobacteria and Bacteroidetes. We observed that positive interactions between nodes indicated niche overlap, while negative interactions indicated competition or variation [67]. In this study, phylogenetically related microorganisms forms well-differentiated clusters (Fig6), and clusters with close correlations among key genera was mainly composed of positive correlation. These results are similar to the co-occurrence network of natural and agricultural soils [66]. In the bacterial network of EM-treated soil, the number of keystone genera and clusters were generally greater than other treatments. In BS treatment, despite a substantial increase in relative abundance of *Bacillus*, it did not become a keystone genus in the microbial network, which further confirmed the previous hypothesis. However, whether these clusters constructed around key genera represent different functional groups remains obscure.

However, the ecological effects of these soil amendments on strawberry cultivation needs a comprehensive evaluation, including the determination of strawberry growth, production and quality in different treatments, and even its long-term effects [10, 68]. At the same time, we will consider the response of a broader range of soils with different physicochemical properties, climate types and field management practices to soil amendments [7, 69]. EM and BS based studies have revealed the effects of soil amendments on bacterial community structure and symbiotic network in strawberry soil. However, the molecular mechanism, phenotypic characteristics, and interactions behind these changes and their effects on plant health remain unclear. Therefore, the q-PCR technique should be used to study how the absolute number of target microorganisms react to soil amendments in agricultural soils. Further metagenomic studies are needed to the accurate determine the beneficial bacteria and pathogens at species level.

## 5. Conclusion

In summary, our research showed that EM2/BS1-treated soil amendments significantly increased bacterial diversity, whereas they had no significant effect on fungal diversity. The effect of the four soil amendments on soil microbiome structure was significant, as all of them reduced the relative abundance of fungal pathogens including *Rhizopus*, *Penicillium* and *Fusarium*. FUNGuild predicted that soil amendments significantly reduced some detrimental functions of soil microhabitat systems (Plant_Pathogen, Animal_Pathogen). Besides, the effects of soil amendments on soil microbial community are mainly indirectly driven by TK, AP and TN, suggesting that the application of soil amendments could have an indirect effect on the soil microbial community by changing environmental factors. Moreover, all soil amendments enhanced the connectivity of bacterial networks, which was the most complex and balanced in EM2-treated soils. Therefore, EM2 and BS1, as novel soil amendments, have the potential to regulate soil microbial community and promote agricultural sustainable development.

## Author contributions

HC, YBL and SLL designed experiments; SLL, MHK, SH, ZYY and YBL carried out experiments; SLL and SH contributed to the preparation of the manuscript and data analyses. HC and YBL supervised the entire study.

## Acknowledgments

This research was supported by the National Natural Science Foundation of China (grant no. 41371262). Meanwhile, we thank the Zhenjiang Institute of Agricultural Sciences for assistance in conduct of strawberry greenhouse trials. We thank Novogene Genomics Institute (Beijing, China) for assistance in bioinformatics analysis.

